# Foreign RNA spike-ins enable accurate allele-specific expression analysis at scale

**DOI:** 10.1101/2023.02.11.528027

**Authors:** Asia Mendelevich, Saumya Gupta, Aleksei Pakharev, Athanasios Teodosiadis, Andrey A. Mironov, Alexander A. Gimelbrant

## Abstract

**Motivation:** Analysis of allele-specific expression is strongly affected by the technical noise present in RNA-seq experiments. Previously, we showed that technical replicates can be used for precise estimates of this noise, and we provided a tool for correction of technical noise in allele-specific expression analysis. This approach is very accurate but costly due to the need for two or more replicates of each library. Here, we develop a spike-in approach that is highly accurate at only a small fraction of the cost.

**Results:** We show that a distinct RNA added as a spike-in before library preparation reflects technical noise of the whole library and can be used in large batches of samples. We experimentally demonstrate the effectiveness of this approach using combinations of RNA from species distinguishable by alignment, namely, mouse, human, and *C.elegans*. Our new approach, `controlFreq`, enables highly accurate and computationally efficient analysis of allele-specific expression in (and between) arbitrarily large studies at an overall cost increase of ~ 5%.

**Availability:** Analysis pipeline for this approach is available at GitHub as `R` package `controlFreq` (github.com/gimelbrantlab/controlFreq).

## 1 Introduction

A highly informative approach to understanding gene regulation is allelespecific expression analysis in RNA-seq data from diploid organisms, including humans. The two parental alleles share the environment of the same nucleus, making allelic imbalance (AI) in gene expression reveal cis-regulation (Nica and Dermitzakis, 2013; Wittkopp and Kalay, 2011). AI analysis has been increasingly used for understanding regulatory variation (Uechi *et al.*, 2020; GTEx_Consortium, 2017; Pirinen *et al.*, 2015; Moyerbrailean *et al.*, 2016; Mohammadi *et al.*, 2019; Vierstra *et al.*, 2020). Besides uncovering genetic effects, allele-specific analysis can reveal epigenetic gene regulation in *cis*, including imprinting (Tucci *et al.*, 2019), X-chromosome inactivation (Galupa and Heard, 2018), and autosomal monoallelic expression (Gimelbrant *et al.*, 2007; Vinogradova *et al.*, 2019; Chess, 2016; Zwemer *et al.*, 2012; Gendrel *et al.*, 2014).

In a previous work (Mendelevich *et al.*, 2021), we have shown that substantial technical noise is present in RNA-seq experiments and significantly varies between experiments, but it is often not accounted for in the analysis. We demonstrated that in the absence of technical replicates, it is impossible to separate this technical noise from biological variation. The resulting underestimation of the technical noise leads to an increased number of false positives in allelic imbalance calls and limited reproducibility between different studies (**Fig.1a**). To quantify and account for this technical overdispersion (**Fig.1b**), we have developed a statistical framework and software tool, `Qllelic`, that takes advantage of the production of two or more RNA-seq libraries for each RNA sample, i.e., technical replicates (Mendelevich *et al.*, 2021). However, preparation and sequencing of replicate libraries at least doubles the experiment’s costs. This complicates practical application of this approach, especially for large projects, where cost of RNA-seq is a major component of the cost of the study.

**Fig. 1.**
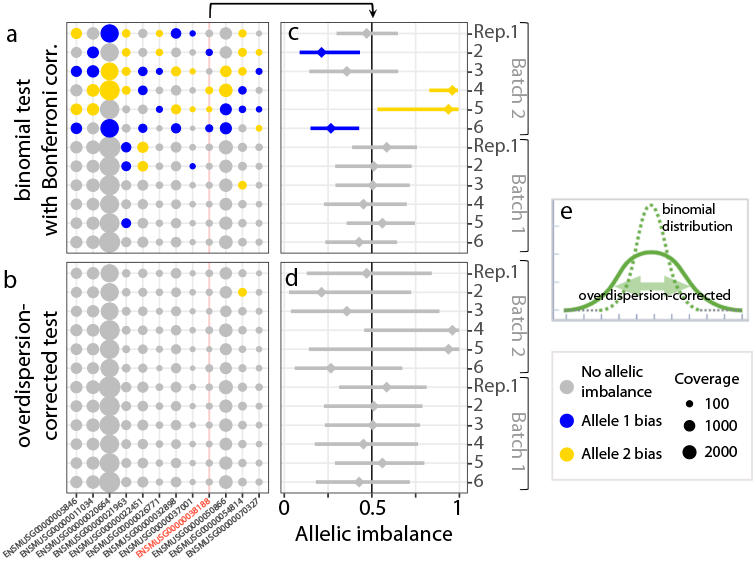
Allele-specific signal from RNA-seq can substantially vary dependent on technical noise. Analysis of replicates can be leveraged to account for this experiment-specific overdispersion. RNA-seq data: 12 technical replicates from the same RNA (129xCastF1 mouse kidney; from (Mendelevich et al., 2021)). “Batch 1” and “Batch 2” were denoted Experiment 2 and Experiment 3 in that paper; both used the same library preparation kit (SMART-Seq v4 Ultra Low Input RNA Kit) with amounts of input total RNA bracketing recommended range: 10 ng per library for Batch 1, and 100 pg for Batch 2. (a) Shown are all genes (coverage >10 allelic counts) with conflicting assignment of allelic bias across 12 RNA-seq libraries from the same RNA. Grey - no allelic imbalance (AI), i.e. null hypothesis of AI=0.5 cannot be rejected at p<0.5 in the binomial test with Bonferroni correction. If null hypothesis rejected (i.e., gene is called as having allelic imbalance): blue - significant prevalence of paternal allele; yellow - significant prevalence of maternal allele. Marker sizes reflect coverage (in total allelic counts); Allelic counts and statistical analysis performed using commonly used approach (e.g., (GTEx_Consortium, 2017)); (b) Same data as in panel (a), with allelic counts table analyzed using modified binomial test with overdispersion correction (AI overdispersion discussed in (Mendelevich et al., 2021)); (c-d) Allelic imbalance for one of the genes (ENSMUSG00000038188, mean coverage of 110 ± 45 allelic counts) denoted in red in (a-b). AI is defined as (maternal counts)/(sum of maternal and paternal counts), so that AI=0 is maternal expression only, AI=1 is paternal only, and AI=0.5 is exactly biallelic. Diamond marker denotes the point estimate for this gene in each replicate library, which is by definition unchanged between (c) and (d). Confidence interval is determined by the statistical test used. Note overdispersion correction (panels b, d) has much more pronounced effect on confidence intervals in replicates from the more noisy datasets (Batch 2) compared to less noisy datasets (Batch 1); (e) Schematic representation of super-binomial overdispersion (solid line) as compared to binomial (dashed line).

Here, we describe an alternative approach for precise allele-specific analysis of large-scale RNA sequencing experiments involving only a small increase in costs. The experimental component of this approach consists of adding an aliquot of foreign, easily distinguishable RNA to every sample (e.g., RNA from heterozygous mouse to human RNA samples) as a spike-in before starting library construction.

Non-allelic RNA spike-in standards have been previously used to access technical noise in transcription levels measurements in bulk RNA-seq (Lovén *et al.*, 2012; Jiang *et al.*, 2011), and estimate background noise, technical batch effects, and doublets in scRNA-seq experiments (Grün *et al.*, 2014; Brennecke *et al.*, 2013; Kim *et al.*, 2015). However, estimates of non-allelic abundance variation are of limited utility for allelespecific analysis (Mendelevich *et al.*, 2021); thus, we aimed to develop an RNA spike-in approach that captures technical noise in allelic imbalance measurements.

The analytical part of our approach relies on the uniformity of the spikein RNA aliquots across all libraries. We extend our previously described idea of processing data from replicates to the spike-in portions of all the libraries in a batch. Crucially, we experimentally show that allele-specific noise properties of the spike-in portion reflect technical noise of the whole sample. Thus, spike-in acts as the standard for estimating technical noise in each library. Adding a fraction of spike-in RNA to the main sample of interest (e.g., mouse RNA to human RNA at 1:10 ratio) removes the need for additional libraries, while only marginally increasing total depth of sequencing, as opposed to doubling or tripling experimental costs.

## 2 Materials and methods

### 2.1 Measuring overdispersion using extended beta-binomial model

In this paper we revised and expanded the procedure described in (Mendelevich *et al.*, 2021). We recall that we define technical replicates as a set of separate libraries prepared from the same RNA, and thus capturing technical noise that accumulates starting from library preparation. Having several technical replicates, we can calculate pairwise Quality Correction Coefficients (QCCs), which one may consider as a widening coefficient of the expected binomial quantiles, for the replicates with similar library sizes. We made pairwise comparisons, and assigned an averaged overdispersion value to each replicate, considering QCC results for relevant pairs.

Here we describe the new procedure, which operates with lots of samples at once (**Fig. 2a**) and assigns individual values of overdispersion measure, iQCC, to each sample in the batch (**Fig. 3a**). It might be applied in both scenarios: spike-ins use case and technical replication use case (see the respective subsections at the end of this section).

**Fig. 2.**
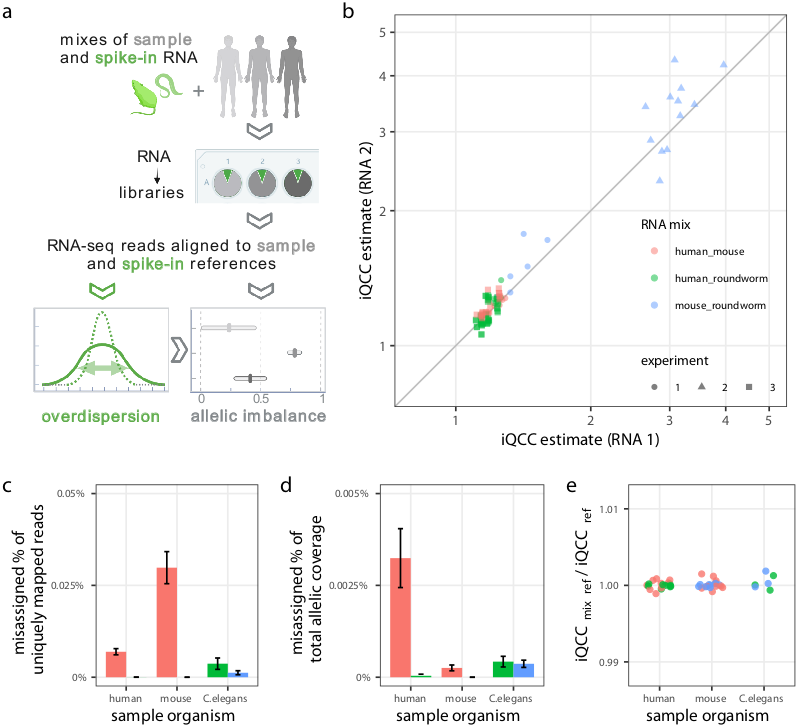
In libraries using mixes of RNA from two distinct organisms, both RNAs show similar allelic overdispersion. (a) Diagram of experimental and computational steps in AI estimation pipeline. Spike-in RNA (green) is added to the main sample (grey) prior to starting preparation of RNA-seq library. (b) Within an RNA-seq library, overdispersion statistic iQCC measured for the two distinct RNAs in a mix is highly similar (Pearson correlation 0.97). For description of libraries in Experiments 1-3, see Table 1. (c-e) Estimation of data loss due to cross-species alignment in data from mixedRNA-seq libraries. Data used: 3 human and 3 mouse biological replicates and 1 roundworm biological replicate (3 technical replicates each), from Experiment 1, aligned on the individual or chimeric reference. (c) Percentage of misaligned reads to the wrong organism among all uniquely aligned reads. (d) Percentage of allelicaly-resolved counts aligned to the wrong reference. (e) Comparison of iQCC values calculated based on alignment to a single or to a mixed (chimeric) reference. (Note also that 0 genes with differential AI were found for each of possible pairs).

**Fig. 3.**
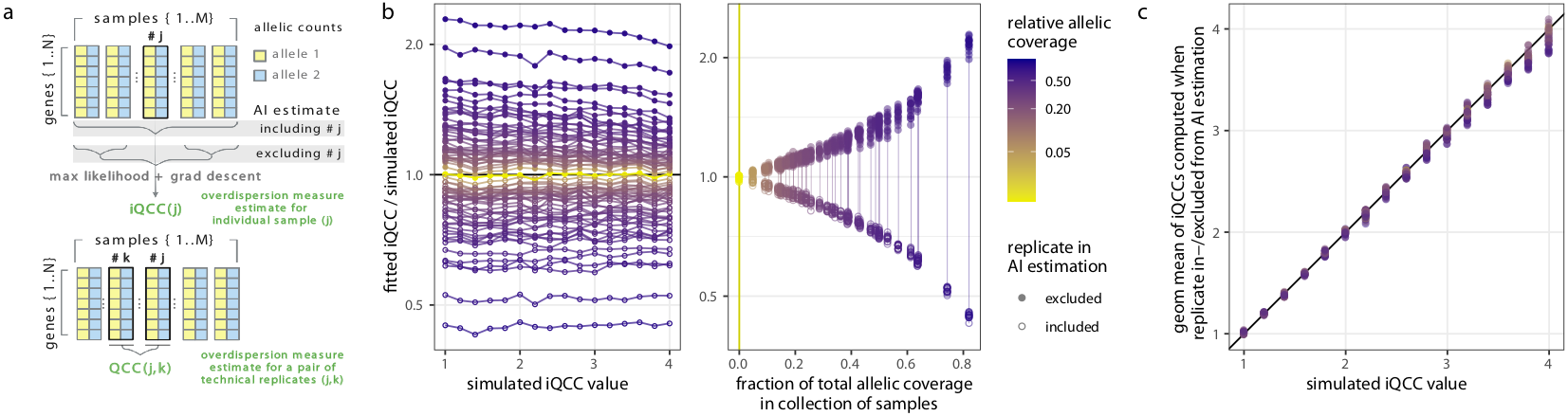
Calculation of iQCC is robust to the number of replicates (spike-ins) and to variation in library sizes within a batch. (a) Scheme of the iQCC calculation (using `controlFreq`) and previously described QCC calculation (using `Qllelic`) for a j-th sample in a batch. See Methods for details. (b) exclusion and inclusion of j-th replicate result in, respectively, under- and overestimation of iQCC from the total pool of replicates. Left panel - as allelic coverage increases, the upper and lower iQCCs estimates converge to ground truth (gold); right panel - extension of panel (a) shows symmetry for overestimated and underestimated measures (lines connect same replicate combinations). (c) same data as in (b), geometric means of underestimated and overestimated iQCC values clearly correlate with the generated levels of overdispersion. (b-c) Simulations of replicate data (all generative iQCCs were similar for all replicates, which mimics technical replication, or similar experimental environment) with different total coverage and overdispersion levels. For iQCC values between 1 (no overdispersion) and 4 (high overdispersion), gene allelic counts for 10 replicates (“libraries”) were generated using predetermined “ground truth” AI distribution and coefficients for total allelic coverage; the procedure was repeated thrice for every set of parameters. Upper and lower iQCC estimates were computed for each selected replicate.

#### 2.1.1 Extended beta-binomial distribution

Beta-binomial family of distributions is commonly used to model an “overdispersed” binomial distribution. One way to parameterize this family is using the real parameters *α* ≥ 0 and *β* ≥ 0 which stand for the number of different balls in the Pólya urn model. Using this parameterization, the probability mass function of the beta-binomial distribution can be written as follows:

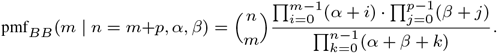

The variance of this distribution is

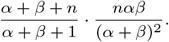

The second part is the variance of the binomial distribution with 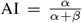, so we capture the overdispersion 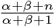 in a parameter we denote by *Q*. Given parameters 0 ≤ AI ≤ 1 and 1 ≤ *Q* ≤ *n*, we can reconstruct back *α* and *β* as 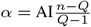 and 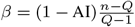.

Two edge cases of the possibility range for *Q* are 1 and *n*. When *Q* approaches *n*, *α* and *β* become close to 0, and the probabilities of all outcomes except for 0 and *n* tend to 0. The limit distribution assigns probability AI to 0, and probability AI to *n*. When *Q* approaches 1, *α* and *β* tend to +∞, and the distribution approaches the binomial distribution with the probability AI.

One drawback of the beta-binomial distribution family is that it doesn’t allow us to model “underdispersed” distributions, in other words, the distributions with *Q* < 1. While most samples give overdispersed distributions, some could be modelled better with underdispersed ones (see Subsection 2.1.4 for more details), and clamping them to one point *Q* = 1 doesn’t make computational sense. Fortunately, we can overcome the problem by extending the probability mass formulas past the edge *Q* = 1, essentially extrapolating the family of beta-binomial distributions into hypergeometric distributions following (Prentice, 1986).

Consider the following modified Pólya urn model with the usual parameters *α* and *β* and an additional parameter *d*. Place *α* ≥ 0 balls of the first type and *β* ≥ 0 balls of the second type into the urn. Draw *n* balls from the urn as usual. The modification is how we replenish the urn. In the beta-binomial case, after drawing a ball we put 2 balls of the same type in the urn, making the total number of balls in the urn larger by 1 on each step. In our modified case, we put *d* +1 balls instead of 2, increasing the total by *d* each time. The probability mass function of this extended beta-binomial distribution is quite similar to the usual one, except that the increment in the multipliers is d instead of 1:

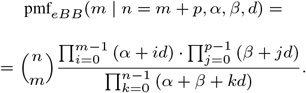

We can see a few properties of this definition right away.

1. The extended distribution doesn’t depend on the scaling of the parameters, i.e. for any *x* > 0

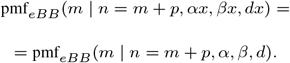
2. If *d* = 1, the extended distribution coincides with the beta-binomial distribution with the parameters *α* and *β*.
3. If *d* = 0, the extended distribution coincides with the binomial distribution with 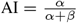.
4. A hypergeometric distribution pmf_*HG*_(*m* | *n* = *m* + *p*, *M*, *P*) belongs to this family as

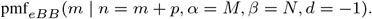

Therefore, this continuous family of distributions contains all betabinomial, binomial, and hypergeometric distributions. From the property 1 we gather that the family is two-dimensional rather than three-dimensional, and can be described using the parameters 0 ≤ AI ≤ 1 and *Q* ≤ *n* just by analytically continuing the description of the beta-binomial distribution. The reparametrization using *AI* and *Q* can be put in the following neat form:

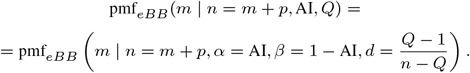

The binomial distribution with *Q* = 1 stops being an edge case, since the range *Q* < 1 becomes treatable, and now we can model underdispersed distributions alongside with the overdispersed ones.

#### 2.1.2 Finding *Q* which maximizes likelihood

For the purpose of using this distribution efficiently in our models, we implement fast and accurate calculation of the density function pmf_*eBB*_(*m* | *n*, AI, *Q*) as follows.

The defining formula of the density function of the extended betabinomial distribution is not suitable for the numerical application right away. Both the numerator and the denominator are orders of magnitude larger than the actual value of the expression, so a straightforward implementation would lose precision. To avoid that, we match each multiplier in the numerator with a suitable multiplier in the denominator, calculate their ratio, and only then take the product. Also, since we are usually interested in the log-likelihood function, we employ one more trick to increase the accuracy of the calculations. Instead of calculating the logarithm of each fraction directly, we calculate it using the function log(1 + *x*). In R language, this function is called `log1p`. Overall, we have

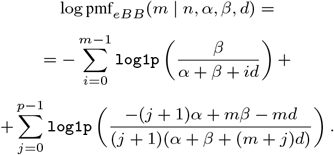

The differential of this expression is straightforward.

These calculations constitute a major building block in the optimization problem which is at the heart of the top-level fitting procedures.

**Given** a vector of allele counts *m_l_*, *p_l_*, 0 ≤ *l* ≤ *g* – 1,
and a vector of corresponding AI estimations AI_*l*_, 0 ≤ *l* ≤ *g* – 1,
**Find** *Q* which maximizes the likelihood function

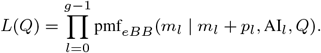

We tackle the problem using the gradient ascent on the logarithm of the likelihood function. The calculation of 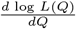 is straightforward and can be easily translated into code. The rest of the implementation comes down to the “flavours” of the gradient ascent.

#### 2.1.3 Computing overdispersion with spike-ins

One of the most important features of the spike-in protocol is a large number of samples involved in overdispersion calculation. This implies that the total allelic counts in spike-in batch are expected to be much higher than in each of the samples. Modelling of the allelic counts in this case is challenging, since in general the sum of two beta-binomial random variables does not need to be beta-binomial. To simplify, we make an assumption that the overdispersion coefficients are approximately the same, in which case the sum will still be beta-binomial, so *m*_1_ + *m*_2_ ~ *eBB*(*n*_1_ + *n*_2_, AI, *Q* = iQCC^2^) when *Q*_1_ ≃ *Q*_2_, *m_i_* stand for the maternal counts, and *n_i_* stand for the total counts. Spike-in samples do not meet technical replication criteria, but they share RNA source. Then the best estimate of AI might be computed from the whole spike-in batch:

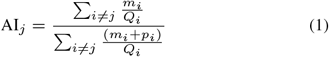

for a selected sample *j* and a set of ground truth *Q*_*i*≠*j*_ values. This estimate can be considered coming from a beta-binomial distribution with *N* being the total coverage of the other samples. When *N* is of several orders of magnitude larger than *n*, the estimate of AI_*j*_ does have negligible variance, and the procedure for fitting the best 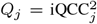 will work well. However, when *n* and *N* are of the same order or *N* is less than *n*, the estimate may deviate from the real AI_*j*_, and the value of *Q_j_* will inevitably be overestimated. This can be particularly challenging if the amount of samples is low. But usually in the spike-in case we have *N* ≫ *n*. See **Fig.3a,b** for how much *Q_j_* will be overestimated depending on the ratio *n*/(*n* + *N*).

If all of *Q_i_* values in a set of samples or *N* ≫ *n* are of similar order, it should be enough to run the fitting algorithm once. The first run of the algorithm sets all *Q_i_* to the same value (*Q_i_* = 1). Thus (1) simplifies to AI_*j*_ = Σ_*i*≠*j*_ *m_i_*/Σ_*i*≠*j*_ (*m_i_* + *p_i_*).

Otherwise, if *Q_i_* estimates from the first run show significant dispersion, the fitting algorithm can be iterated. In the grand scheme, we alternate between fitting *Q*’s, then fitting AI’s, and so on. The AI fitting is run using the formula (1), and the *Q* fitting is run using the gradient ascend algorithm. After each *Q*-fitting step, *Q_i_* values are compared to previous estimates, and iterations are repeated until convergence.

#### 2.1.4 Computing overdispersion in case of technical replication

Here we need to take care of the overestimation of *Q_j_*. To do that, we counter the previous estimate with a different one. If one includes the sample *j* in the AI_*j*_ estimation, then the corresponding fitted *Q_j_* will be underestimated. Moreover, on the simulated data we see that the over- and underestimated iQCC values deviate from the real value by *the same amount* in the logarithmic scale, see **Fig.3b**. We use this fact in the case of technical replication, where we fit iQCC_*j*_ to be the geometrical mean of the overestimate and the underestimate to get a more precise statistic (**Fig.3c, Suppl.Fig.6**). Using this more precise fitting, we can safely use the pipeline on a small amounts of samples, given a prior knowledge that the samples have similar overdispersion.

### 2.2 Data

Information on RNA-seq datasets generated for this work, including sources of RNA, library preparation methods, sequencing, and data processing is summarized in **Table 1**.

**Table 1.**
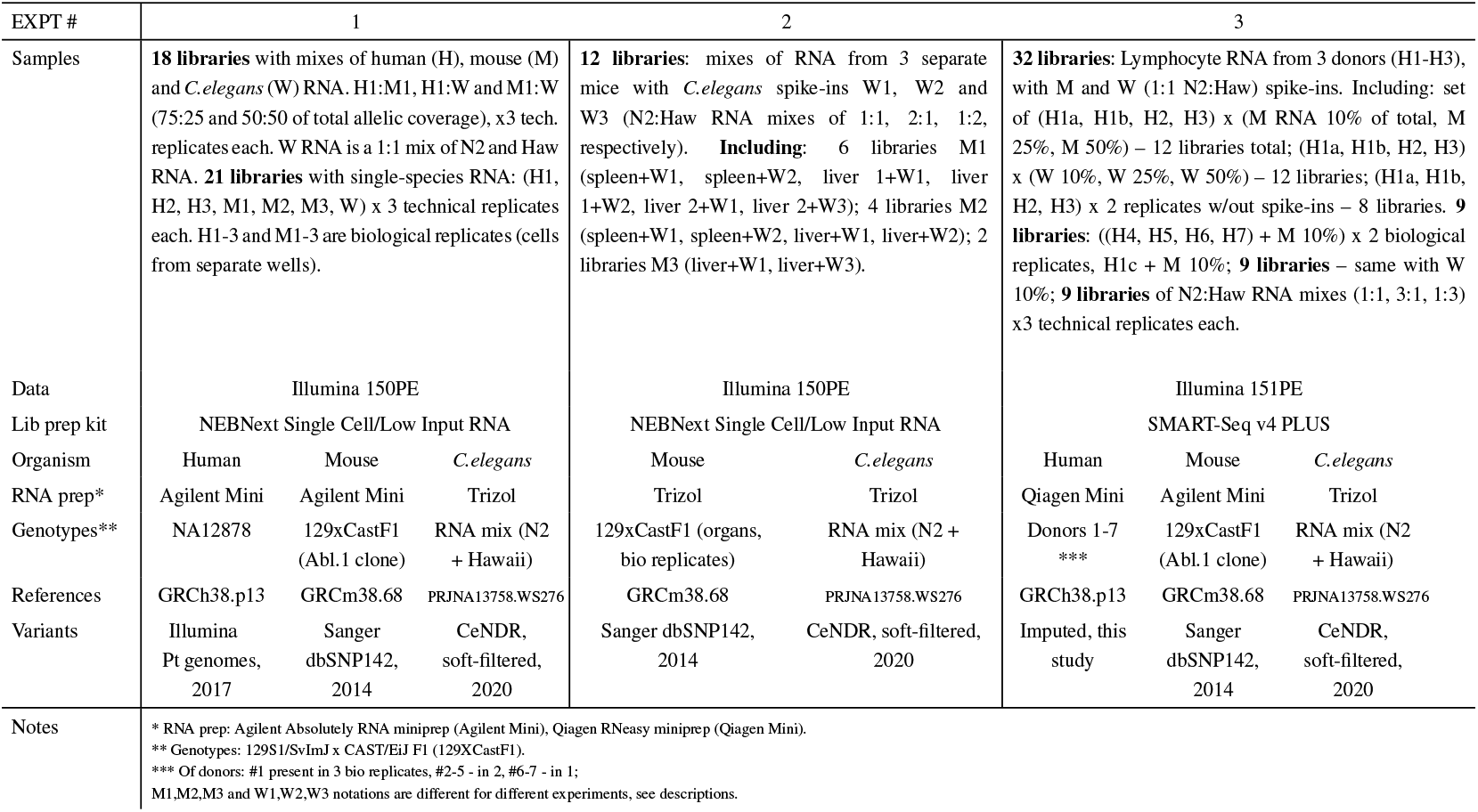
Description of the data used in this work.

#### Biological material

v-Abl pro-B clonal cell lines Abl.1 derived previously from 129S1/SvImJ × Cast/EiJ F1 female mice (Zwemer *et al.*, 2012) and human clonal pro-B cell line GM12878.DF1 (Nag *et al.*, 2013) were cultured in RPMI medium (Gibco), containing 15% FBS (Sigma), 1X L-Glutamine (Gibco), 1X Penicillin/Streptomycin (Gibco) and 0.1% *β*-mercaptoethanol (Sigma). Human peripheral blood monocytes from deidentified donors were appropriately consented for generating genomic sequencing data for unrestricted deposition in public databases. *C. elegans* strains N2 and CB4856 (Hawaiian) were maintained at 20°C on nematode growth medium (NGM) plates with Escherichia coli OP50 bacteria (Brenner, 1974).

#### RNA preparation

Biological replicates for human and mouse clones (see **Table 1**) were made by growing cells in separate wells of a 6-well plate at a seeding density of 500,000 per mL (1,500,000 total cell per well). RNA isolation and DNase treatment for both clonal cell lines was performed using Absolutely RNA Microprep Kit (Agilent) according to instruction. RNA from C.elegans was prepared using Trizol reagent (Invitrogen), with DNase treatment using TURBO DNA-free kit (Ambion). RNA integrity was assessed using Bioanalyzer and was quantified using Qubit RNA HS Assay. DNase-treated RNA from N2 and Hawaiian strains was mixed in proportions described in **Table 1**. For RNA preparation from mouse tissues, whole tissues were collected from adult 129xCastF1 mice housed at the DFCI mouse facility, with parent animals obtained from the Jackson Laboratories. All animal work was performed under DFCI protocol 09-065, approved by the DFCI Institutional Animal Care and Use Committee. Animals were housed in accordance with Guide for the Care and Use of Laboratory Animals. Collected tissues were crushed using mortar and pestle in liquid nitrogen. This powdered tissue was either taken directly for RNA isolation using Trizol reagent or stored in liquid nitrogen for later use.

#### Library preparation and sequencing

For experiments 1 and 2 (see **Table 1**), aliquots of total RNA were used to prepare libraries using NEBNext Single Cell/Low Input RNA Library Prep Kit (NEB). The libraries were sequenced at Novogene on the Illumina NovaSeq platform and 150 bp paired-end reads were generated. For experiment 3 (see **Table 1**), libraries were prepared using SMART-Seq v4 PLUS kit (Takara), and PE-151 sequencing was performed on Illumina NovaSeq at the High Scale Data Unit at Altius Instutite for Biomedical Sciences.

#### Variant calling and imputation

Imputation on genotype calls for donor samples in experiment #3 was performed on autosomes only. VCFTools was used to split files per chromosome. Per chromosome genotype files were submitted to the TopMed imputation server (imputation.biodatacatalyst.nhlbi.nih.gov) with the following parameters: imputation=minimac4-1.0.2; phasing=eagle-2.4; panel=TOPMED.vR2; Mode: Quality Control & Imputation. 131,562 sites were excluded prior to imputation, leaving 572,955 sites for imputation. 131,452 of the excluded sites were due to monomorphic sites. After imputation, only markers with a R2 value of greater than .9 were kept. *De novo* SNPs were annotated with dbSNP v151.

### 2.3 Generation of allelic count tables

#### Reference preparation

To receive *in silico* chimeric pseudogenome reference files “containing” more than one organism, we first create a set of respective pseudogenome reference files for each of them, namely, pseudoreference fasta files for both alleles (reference genome with inserted SNPs related to one fixed haplotyped allele), allelic VCF files (SNPs only, and 1st and 2nd alleles playing role of reference and alternative respectively) and reference GTF files. The procedure is described in detail in (Mendelevich *et al.*, 2021). Next, we merge them with a consistent renaming of similar chromosome names via appending the distinguishing suffixes (for example, chromosome 1 may be renamed to “1m” in mouse and “1h” in human These resulting files are used as “reference” files in all the next steps. Finally we index an *in silico* chimeric pseudogenome with `STAR`, setting `sjdbOverhang` parameter to the advised data-specific `len(read)-1`.

#### Data pre-processing

Note that we have revised recommendations on allelic counts tables creation relative to Qllelic, to minimize effect of noise coming from statistically dependent counts for nearby variants while preprocessing RNA-seq data (see GitHub for details: github.com/gimelbrantlab/controlFreq). The unit of overdispersion analysis is a table with allelic read counts per gene. In order to obtain that kind of tables we first align the reads to both pseudogenomes with STAR (v2.7.9a), which supports allele-aware alignment (`–outSAMattributes vA vG`) when provided prearranged reference fasta and vcf (see section above for details). For further analysis we select reads aligned at positions that include at least one SNP to determine the allele, and filter out those whose SNP signals or alignment positioning on two haplotypes don’t show consistency. The remaining reads are assigned to one of the alleles, according to SNP calls and alignment quality. Finally, we assign allele-resolved reads to genes and calculate gene counts using `featureCount` (v2.0.2) (with parameters `–countReadPairs –B –C`). Autosomal genes were used for iQCC values calculation.

## 3 Results

### 3.1 Mixes of RNA from highly distinct organisms show similar overdispersion in all components

We showed previously (Mendelevich *et al.*, 2021) that AI overdispersion in poly-A RNA-seq experiments behaves like a consistent multiplicative parameter from binomial expectations, across all genes in a sample. We hypothesized that if we mix RNA from two highly distinct species, the overdispersion will be similar for all components. We developed the experimental and computational protocol (operating with evolved overdispersion measure iQCC, individual quality correction coefficient, see **Fig. 2a**), and have shown the proximity of overdispersion measured across different regions of the genome, including chromosomes of different species origins in our *in-silico* chimeras (see **Fig. 2b**, **Suppl.Fig. 5**, **Fig. 4**).

**Fig. 4.**
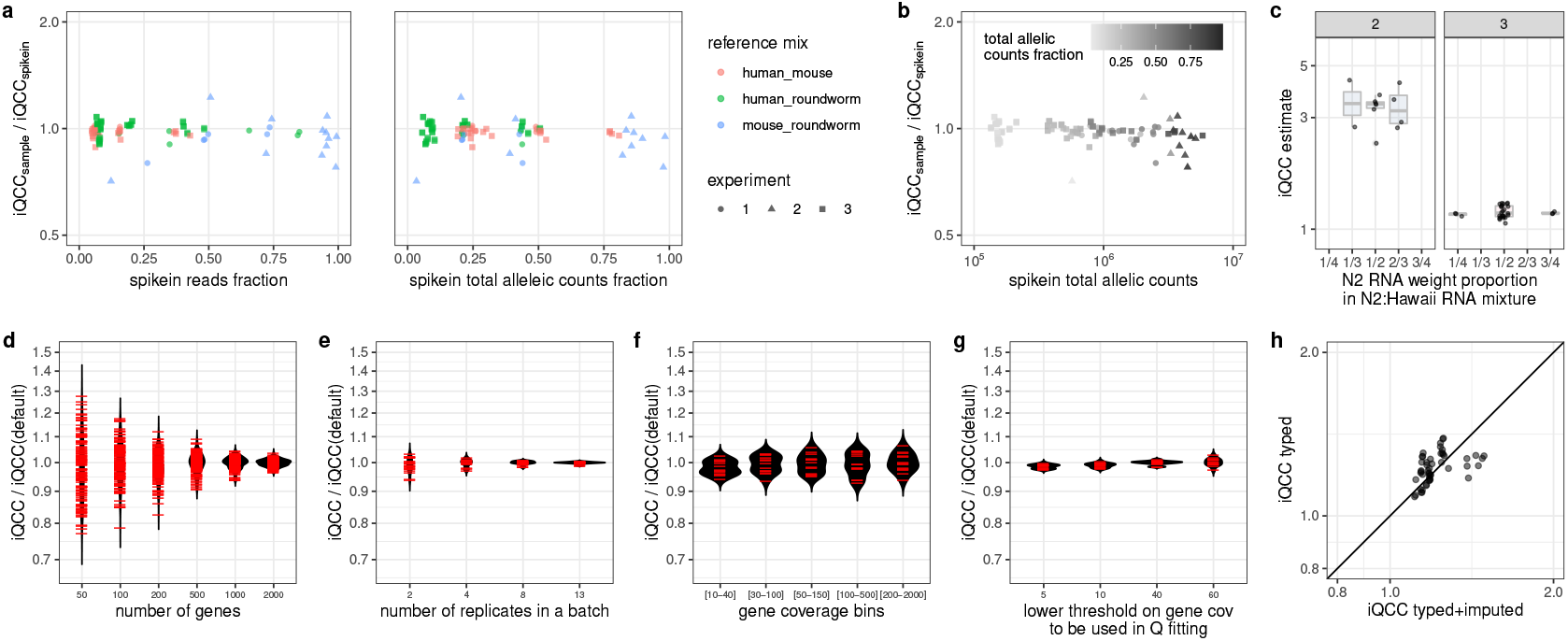
Overdispersion estimate is robust to wide range of variation in the spike-in quantity and composition. (a) Ratio of iQCC values for sub-library from organism 1 and from organism 2 (assigned as “main sample” and spike-in) remains close to 1 with diminishing fraction of spike-in RNA in the initial mix (see Table.1 for details); (b) Ratio of iQCC values for organism 1 and organism 2 for different total allelic counts (reflecting library size) and the spike-in fraction. Note decreased fidelity for extremely low spike-in counts (tens of thousands allelic reads). Note: in the case of equal mix compounds organism 2 is assigned to “spike-in sample”; (c) Polymorphism in the spike-in RNA can be mimicked by a mix of RNA from two distinct strains, with no strict requirement that mix is exactly 1:1. iQCC values are robust across mixes of RNA from N2 and Haw strains of C.elegans with different allelic ratios; (d) iQCC computed on randomly sampled N genes for 13 replicates (10% spike-in samples), shown for each replicate separately. Each gene sampling was performed 10x; (e) iQCC computed on different collections of samples: 11 pairs, 3 quadruples, 3 eights and 2 batches of 13 replicates; (f) iQCC computed in different allelic coverage bins, on 13 replicates (10% spike-in samples), only genes whose coverage belong to the interval in all 13 replicates were considered (number of genes per bins: 611, 610, 399, 496, 232); (g) iQCC computed with different coverage threshold for fitting, using whole data, on 13 replicates (10% spike-in samples); (d-g) Data: experiment 3, mouse spike-ins. Default iQCC values are computed with fitting threshold on coverage = 30 and in a batch of all 21 spike-in samples, without any other restrictions; (h) iQCC computed on experiment #3 samples containing human RNA, when only typed or typed and imputed variants are considered.

Notably, human, mouse and roundworm data can be mapped simultaneously without too much interference (see **Fig. 2c-e**), which makes it possible to use those organisms as spike-in add-ons to each other.

### 3.2 Use of one RNA across many samples can serve as a “stand-in” for tech replicates

Overdispersion measurement analysis on spike-in components differs from analysis of technical replicates, since here we should not be bound by the strict assumption of shared overdispersion between all samples present in a batch. Aiming to meet the changed criteria we had to modify and improve our previous overdispersion calculation pipeline, which currently takes advantage either from same overdispersion (technical replicates case) or from replicates quantity (spike-in case), and utilizes the assumption of shared initial state of allelic proportions in RNA prep in both scenarios (see **Fig. 3a**, and Methods). Finally, overdispersion estimates (iQCC) obtained with the current procedure are highly correlated with `Qllelic` results (see **Suppl.Fig. 5**).

### 3.3 Spike-in protocol is flexible and allows variation of parameters

Aiming to determine the limitations of spike-in methodology, we varied and tested different conditions. Combinations of different organisms pairs, mixed in different proportions, resulted in comparable overdispersion measurements on both components (see **Fig. 4a**). More specific, the only limitation is total allelic coverage (see **Fig. 4b,f**), which clearly depends on the spike-in proportion, total number of sequenced reads and SNP density in an organism (for human it is on the order of 1/1000, for mice and roundworm samples we used it is 1/100 and 1/500 respectively). Moreover, mixes of RNA prepared from individual parental strains of *C.elegans* (N2 and Hawaiian (CB4856)) performed as a spike-in just as well as N2xHawaiian F1 heterozygous cross (see **Fig. 2b** and **Fig. 4a**). We also showed that N2 and Hawaiian RNA could be mixed in a range of proportions with similar overdispersion estimates (see **Fig. 4c** and Discussion). This makes experimental preparation of spike-ins much easier than worm or mouse husbandry.

## 4 Discussion

The spike-in approach we describe here, including the data-processing pipeline, is much more accurate and precise in estimating allele-specific expression than the common single-replicate design (**Fig. 1**). In fact, it is at least as accurate as `Qllelic`, the approach with technical replicates for each sample we described previously, but rather than costing several-fold higher, the cost increases only by ~ 5 – 10%.

The new approach, `controlFreq`, supersedes our previous one, `Qllelic`, and includes its functionality when operating with a small number of samples with the assumption about overdispersion similarity between them (for example, in technical replicates).

The improvements of computation protocol include moving from pairwise overdispersion measure Quality Correction Coefficient, computed by `Qllelic` on pairs of technical replicates, to batch-wide calculation of individual Quality Correction Coefficients (iQCC) for each sample. Besides saving costs, the new approach is superior compared to generating two or three technical replicates since for each sample we assess overdispersion from a large number of spike-in replicates. This enables confident detection of outliers, allows us to remove the assumption of similarity of overdispersion in all samples, and eliminates expectations of library size similarity. Overall, the analysis is highly robust to fluctuations.

An important practical feature of the spike-in approach is that the spike-in RNA must be identical only within the batch of samples being analyzed. Once the overdispersion corrections are calculated for the batch of samples, they can be correctly compared to samples from any other batch, including those using entirely different spike-ins (or, indeed, any sample with properly estimated AI overdispersion). This means that spike in RNA can be prepared as needed, and does not have to come from a single source.

As far as we can establish, the only requirements of the spike-in material are (i) that it has significant density of polymorphisms (the higher the better) and (ii) its nature enables it to undergo the same library preparation process as the main sample. So, for example, bacterial RNA should not be used for poly-A-based libraries, or *C.elegans* RNA as a spike-in standard for mammal samples when using library protocol with ribo-depletion. Otherwise, spike-in material can be produced in any convenient way; we posit that once the “snapshot” of allelic proportions is captured for the whole library, the origin of library components does not matter. For example, we used “synthetic F1s” – mixes of RNA from homozygous *C.elegans* strains (see **Fig.4c**) as a spike-in, which is easier than breeding F1 crosses. It might be possible to use yeast RNA as a spikein, or develop a set of synthetic molecules for allele-resolved RNA-seq analysis, analogous to ERCC controls (Jiang *et al.*, 2011).

We note that the experiments we described are focused on poly-A-enriched bulk RNA sequencing. While we expect that application of this approach can be extended into a variety of other experimental settings where accurate measurement of allelic signal is desirable, a pilot study should be performed in each particular case to test the applicability of spike-in methodology and tune it accordingly. A clear example of an application is capturing technical noise in single-cell RNA-seq; this is a question of great importance since otherwise (Kim *et al.*, 2015), no technical replication is readily available at the single-cell level (attempts to split single-cell RNA into two replicates are extremely technically challenging (Deng *et al.*, 2014)). It is known that the rate of overdispersion in single-cell analysis is high, underscoring the need for reliable methods for distinguishing technical noise from meaningful biological variation. Finally, a similar approach might be applicable to other types of libraries besides RNA, such as ChIP-seq, ATAC-seq and analysis of DNA methylation.

## Acknowledgements

The authors thank Max Heiman and his lab for sharing the *C.elegans*; Andrew Bortvin and Natalia Koutseva for expert technical support; Jess Halow and Sadie Patraw for work with human cells; Dan Bates and High Scale Data Unit at Altius Instutite for sequencing work; A.Bortvin, C.Edwards and members of the Gimelbrant lab for insightful comments. Figures 2a and 3a were created with BioRender.com.

