## Supplementary methods, Supplementary figures (Fig.5 and Fig.6) for "Foreign RNA spike-ins enable accurate allele-specific expression analysis at scale"

### Supplementary information

#### Supplementary methods

**Extended beta-binomial distribution**

A point we should address is whether the formula for the extended beta-binomial distribution gives rise to a proper mass probability function with non-negative values and the total sum 1 of the values. Any parameter $1 < Q < n$ specifies the formula to the probability mass function of a beta-binomial distribution. Since the expression $\mathrm{pmf}_{eBB}(m \mid n = m + p, \mathrm{AI}, Q)$ is an analytic function of AI and $Q$, we gather that the total sum of the probabilities is indeed 1 for any parameters AI and $Q$.

We can also directly access the values of $Q$ such that $\mathrm{pmf}_{eBB}(m \mid n, \mathrm{AI}, Q) \geq 0$ for each $0 \leq m \leq n$. In terms of $\alpha$, $\beta$ and $d$, we need the inequalities

$$\alpha + (n-1)d \geq 0,$$
$$\beta + (n-1)d \geq 0,$$
$$\alpha + \beta + (n-1)d > 0.$$

Note that in terms of the Pólya urn model, the inequalities correspond to the possibility of trying to take a ball of a certain type one more time than the capacity of the urn allows. In the urn corresponding to a hypergeometric distribution, this problem is non-existent: any such scenario has the probability 0. In the extended beta-binomial distribution case, the formulas do not give 0 that easy because $\alpha$ and $\beta$ may not be integer multiples of $d$.

We apply an adhoc trick to match the behaviour to the hypergeometric case more closely. Take a negative $d$ such that the inequality $\alpha + \beta + (n-1)d > 0$ still holds. Then the formulas for $\mathrm{pmf}_{eBB}(m \mid n, \alpha, \beta, d)$ must give a positive value for at least some $0 \leq m \leq n$ (to be precise, at least for $m$ closest to $\frac{n\alpha}{\alpha+\beta}$). On the other hand, the formulas may fail to give positive value for $m$ close to an edge, where one of the inequalities $\alpha + (m-1)d > 0$ or $\beta + (n-m-1)d > 0$ fail. Disregard the value of the formula at these points, and set $\mathrm{pmf}_{eBB}(m \mid n, \alpha, \beta, d)$ as 0 for these $m$. Finally, rescale the whole density function so that the total sum of left positive values is equal to 1.

In this way we have extended the definition of the distribution up to the point $d = -\frac{\alpha+\beta}{n-1}$. In terms of $Q$, the distribution is defined for all $Q > -\frac{1}{n-2}$. One may be surprised by a negative value of $Q$, since we first introduced it as a multiplier in the calculation of the variance. The apparent contradiction is resolved by noting that $Q$ stops to be the precise measure of the overdispersion once we force at least one value of the density function to be 0 instead of the real value of the formula.

**Calculating the pdf of the extended beta-binomial distribution**

Let us present the derivation of the matching here. Without loss of generality, assume that $\alpha \geq \beta$. The matching is as follows:

$$\mathrm{pmf}_{eBB}(m \mid n, \alpha, \beta, d) = \binom{n}{m} \frac{\prod_{i=0}^{m-1} (\alpha + id) \prod_{j=0}^{p-1} (\beta + jd)}{\prod_{k=0}^{n-1} (\alpha + \beta + kd)} =$$
$$= \frac{\prod_{i=0}^{m-1} \frac{\alpha+id}{i+1} \prod_{j=0}^{p-1} \frac{\beta+jd}{j+1}}{\prod_{k=0}^{n-1} \frac{\alpha+\beta+kd}{k+1}} =$$
$$= \frac{\prod_{i=0}^{m-1} \frac{\alpha+id}{i+1}}{\prod_{k=0}^{m-1} \frac{\alpha+\beta+kd}{k+1}} \cdot \frac{\prod_{j=0}^{p-1} \frac{\beta+jd}{j+1}}{\prod_{k=m}^{n-1} \frac{\alpha+\beta+kd}{k+1}} =$$
$$= \prod_{i=0}^{m-1} \frac{\alpha + id}{\alpha + \beta + id} \cdot \prod_{j=0}^{p-1} \frac{m+j+1}{j+1} \frac{\beta + jd}{\alpha + \beta + (m+j)d}.$$

Now the expression is the product of $n$ fractions, with each the fractions having the order much more closer to 1.

With regards to the derivative, we do not need to employ any particular calculation tricks. The final expression, after the substitution of the variables and some algebraic transformations, is as follows.

$$\frac{d \log \mathrm{pmf}_{eBB}(m \mid n, \mathrm{AI}, Q)}{dQ} =$$
$$= \sum_{i=0}^{m-1} \frac{(1-\mathrm{AI})(n-1)i}{\big((i-\mathrm{AI})Q + (\mathrm{AI}n - i)\big)\big((i-1)Q + (n-1)\big)} +$$
$$+ \sum_{j=0}^{p-1} \frac{(n-1)\big(\mathrm{AI}j - (1-\mathrm{AI})m\big)}{\big((j+\mathrm{AI}-1)Q + ((1-\mathrm{AI})n - j)\big)\big((m+j-1)Q + (p-j)\big)}$$

**The gradient descend flavours for finding optimal $Q$**

The most important aspect of the likelihood function that influences the procedure is that it is defined only on a segment from some negative value $Q = Q_{min}$ up to $Q = Q_{max} = \min_l m_l + p_l$.

First, we must choose a way to convert the derivative into the value of an ascend step. To do that, before the ascend we gauge the possible range of the values of the derivative at 5 fixed points in the domain of the likelihood function. We take the median as the estimate for the median value of the derivative, and in the procedure we normalize each derivative by this median value.

To decrease the probability of hopping left and right around a local maximum, or taking painfully long to reach a maximum with a “plateau” around it, we introduce a tracking integer variable `streak`. The ascend step gets multiplied by $1.1^{\texttt{streak}}$. When the derivative is coaligned with the previous step, we increase `streak` by one, and when the derivative is misaligned with it, we decrease `streak` by one. We also reset `streak` to zero before changing its trend.

Two bounds of the domain of the likelihood function are different. The $Q_{min}$ bound is there because the limit of $\log\ L(Q)$ is $-\infty$ as $Q$ tends to $Q_{min}$. It means that the derivative function also becomes more extreme closer to $Q_{min}$. We could counteract this by substituting with a different variable and thus eliminating the extreme behaviour of the derivative, but functionally the same behaviour can be achieved by again adjusting the constant of the derivative-to-step transformation. This time, we multiply the step by $Q - Q_{min}$.

The $Q_{max}$ bound is of a different nature. The limits of both $\log\ L(Q)$ and $\frac{d\ \log\ L(Q)}{dQ}$ exist when $Q$ tends to $Q_{max}$. It could happen that the global likelihood maximum is achieved at $Q = Q_{max}$. In this case, we make a bioinformatics-informed decision to filter out the genes with this coverage, and repeat the ascend again. After filtering out, we can end up with no genes. In this case we return $Q = +\infty$.

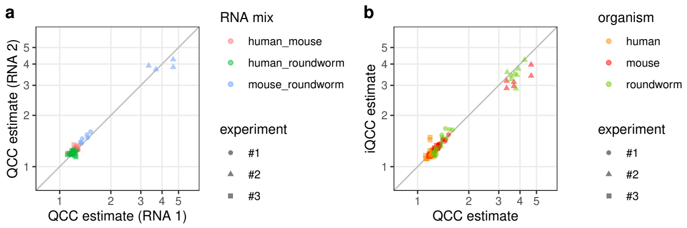


**Fig. 5.** (Supplementary) Independent QCC estimates support our iQCC observations. (a) QCC estimates correlation for 2-component RNA mixes. (Pearson correlation = 0.98). (b) iQCC measurements correlate with QCCs (computed with Qllelic, Pearson correlation 0.98). (a-b) Data: all mixed samples from Experiments 1-3. QCC values were computed for pair of replicates with total allelic counts difference $\leq 15\%$, and geometrical mean of QCC estimates was assigned to a replicate when it was involved in several pairs comparisons.

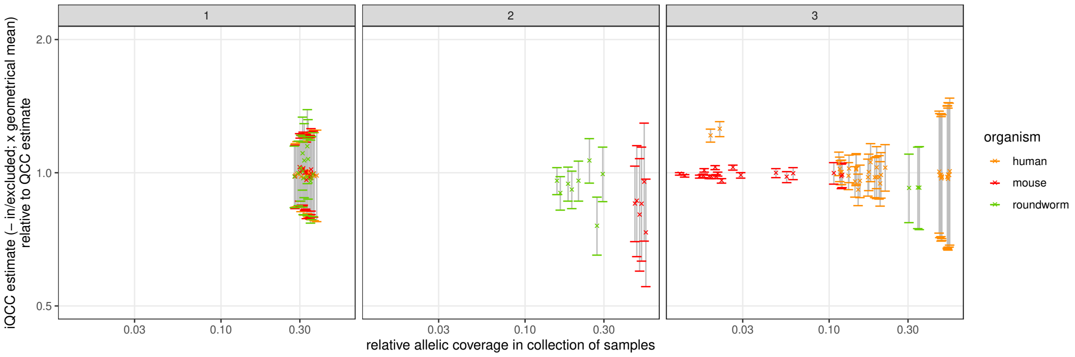


**Fig. 6.** (Supplementary) Convergence of upper and lower iQCC estimates and geometric mean estimate to QCC values computed with `Qllelic` on mixed samples in Experiments 1, 2 and 3 (same data as in Suppl.Fig.5) with increasing of total alellic coverage difference.
